# White LED Lighting for Plants

**DOI:** 10.1101/215095

**Authors:** Anton Sharakshane

## Abstract

The highest intensity of photosynthesis is obtained under red light, but plants die or their growth gets disrupted if only red light is used. For example, Korean researchers [1] have shown that under pure red light the amount of the grown lettuce is greater than under a combination of red and blue light, but the leaves have a significantly smaller amount of chlorophyll, polyphenols and antioxidants. And the researchers at the Faculty of Biology of the Moscow State University [2] have found that the synthesis of sugars is reduced, growth is inhibited and no blossoming occurs in the leaves of Chinese cabbage under narrow-band red and blue light (as compared to a sodium lamp).

What kind of lighting is needed to get a fully developed, large, fragrant and tasty plant with moderate energy consumption?

## How to measure energy efficiency of a plant growth fixture?

Main metrics to measure energy efficiency of phyto light:

- *Photosynthetic Photon Flux* (*PPF*), in micromoles per joule, i.e. in the number of light quanta within the range of 400-700 nm, emitted by a fixture that consumed 1 J of electricity.
- *Yield Photon Flux* (*YPF*), in effective micromoles per joule, i.e. in the number of quanta per 1 J of electricity, taking into account the multiplier – the *McCre*e curve.

*PPF* is always slightly higher than *YPF* (the *McCree* curve is normalized to unity and in the greater part of the range it is less than one), therefore it is advantageous for the sellers of light systems to use the first metric. The second metric is more advantageous for buyers, as it estimates energy efficiency more adequately.

**Fig. 1.**
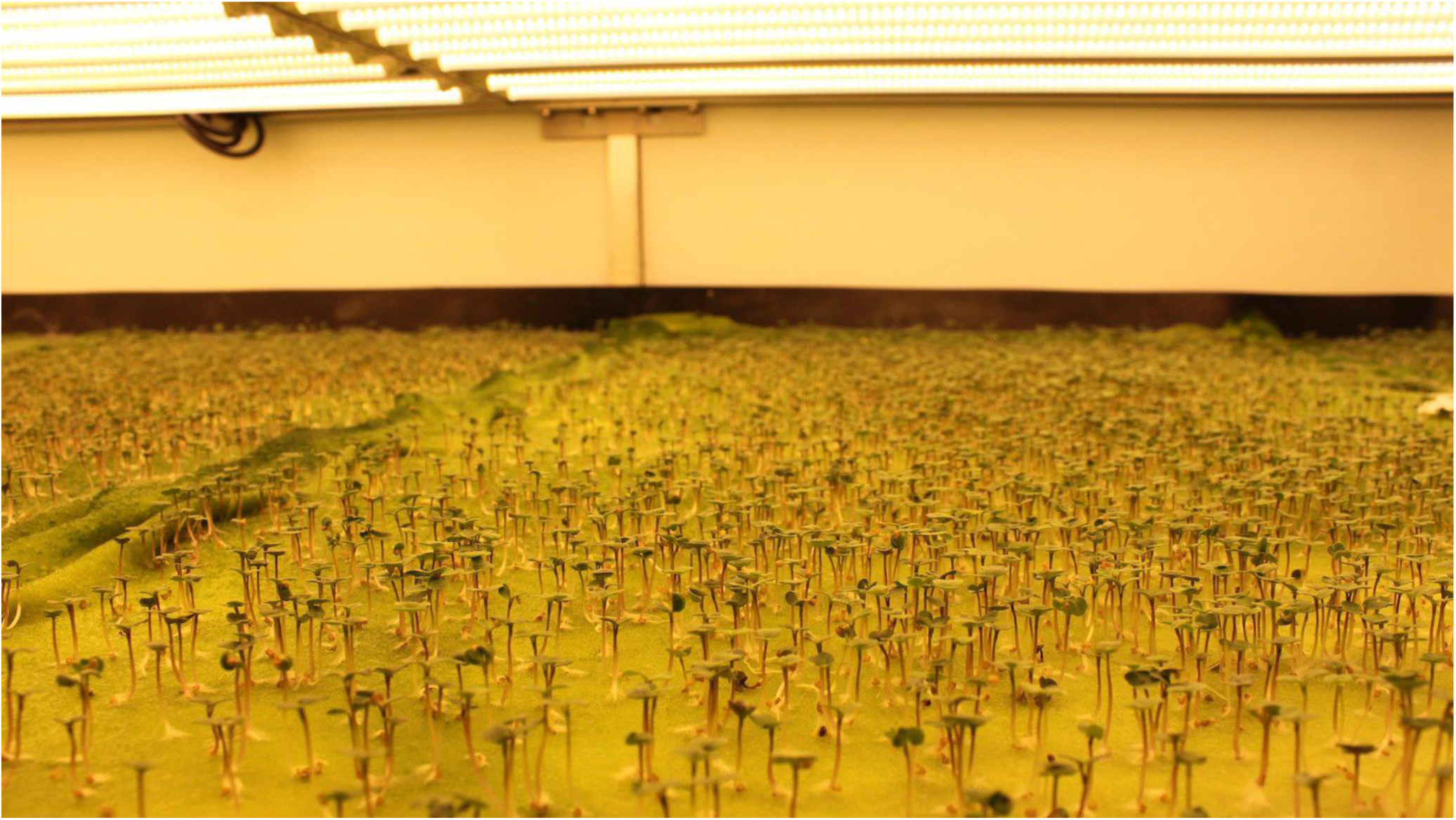
Leanna Garfield, *Tech Insider — Aerofarms*

## Efficiency of HPS lamps

Large farms with vast experience that watch their money still use sodium lamps. Yes, they willingly agree to install LED lamps above their experimental beds, when the lamps are provided for free, but they do not agree to pay for them.

Fig. 2 shows that efficiency of a sodium lamp strongly depends on power and reaches its maximum at 600 W. The characteristic optimistic value of *YPF* for a 600-1,000 W sodium lamp is 1.5 eff. μmol / J. Efficiency of 70-150 W sodium lamps is smaller by half.

**Fig. 2.**
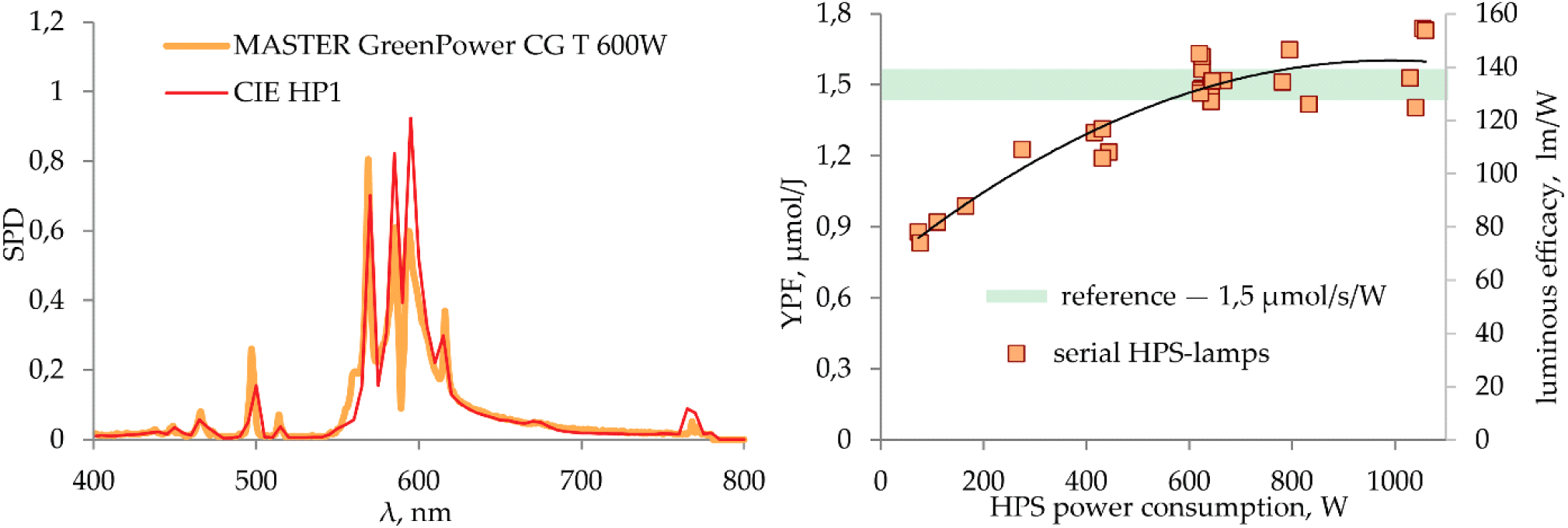
Typical spectrum of a sodium lamp for plants (*left*). Efficiency in lumens per watt, and in effective micromoles of standard sodium lamps *Cavita, E-Papillon, Galad* and *Reflux* for greenhouses (*right*)

Any LED lamp with efficiency of 1.5 eff. μmol / W and an acceptable price can be considered a worthy replacement for a sodium lamp.

## Doubtful efficiency of red and blue phyto lamps

In this article, we do not mention the chlorophyll absorption spectra because it is inappropriate to refer to them in the discussion about using light flux by a living plant. Chlorophyll *in vitro*, isolated and purified, does absorb only red and blue light. In a living cell, pigments absorb light within the entire range of 400-700 nm and transfer its energy to chlorophyll. Energy efficiency of light in a leaf is determined by the *1972 McCree* curve (Fig. 3).

**Fig. 3.**
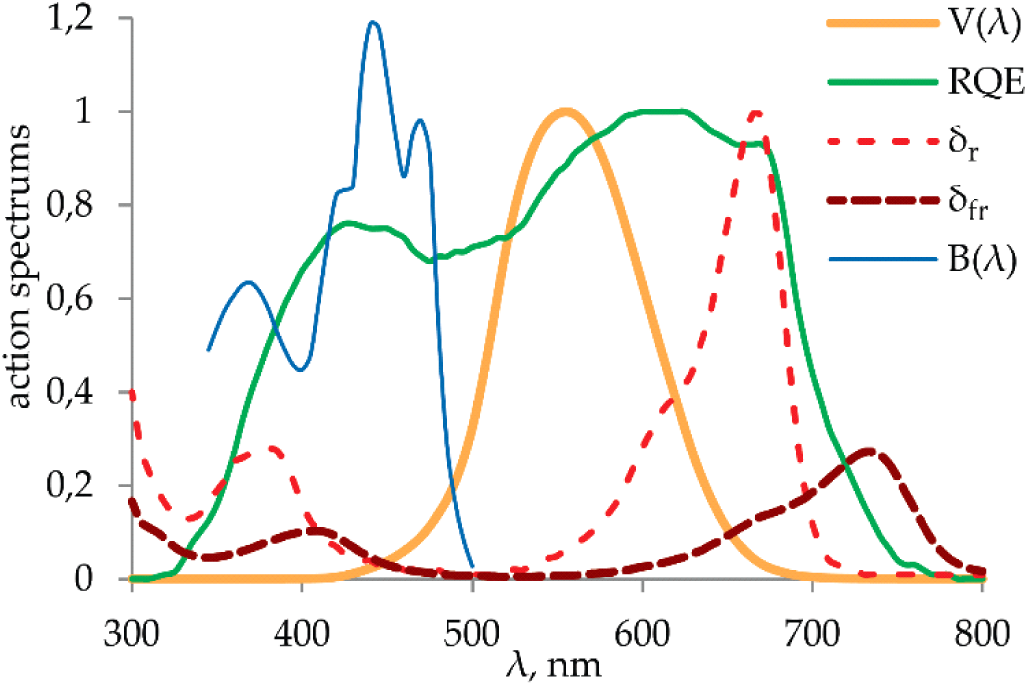
*V*(λ) — human visibility curve; *RQE* — relative quantum efficiency for a plant (*McCree* 1972); *σ*_*r*_ and *σ*_*fr*_ — curves of absorption of red and far red light by phytochrome; *B*(λ) — phototropic efficiency of blue light [3]

Note: Maximum efficiency in the red range is about 50% higher than the minimum efficiency in the green range. And if you average the efficiency over any broad band, the difference becomes even less noticeable. In practice, redistribution of some of the energy from the red to the green range sometimes, on the contrary, strengthens the energy function of light. Green light passes through the leaves to the lower tiers, the effective leaf area of a plant increases sharply, and the yield of, for example, lettuce rises [2].

## White LED lighting for plants

The energy feasibility of plant lighting using common white LED lamps has been studied in the paper [3].

The characteristic shape of the white LED spectrum is determined by:

- Balance of short and long waves correlated with the color temperature (Fig. 4, left);
- Spectrum occupancy degree correlated with the color rendering (Fig. 4, right).

**Fig. 4.**
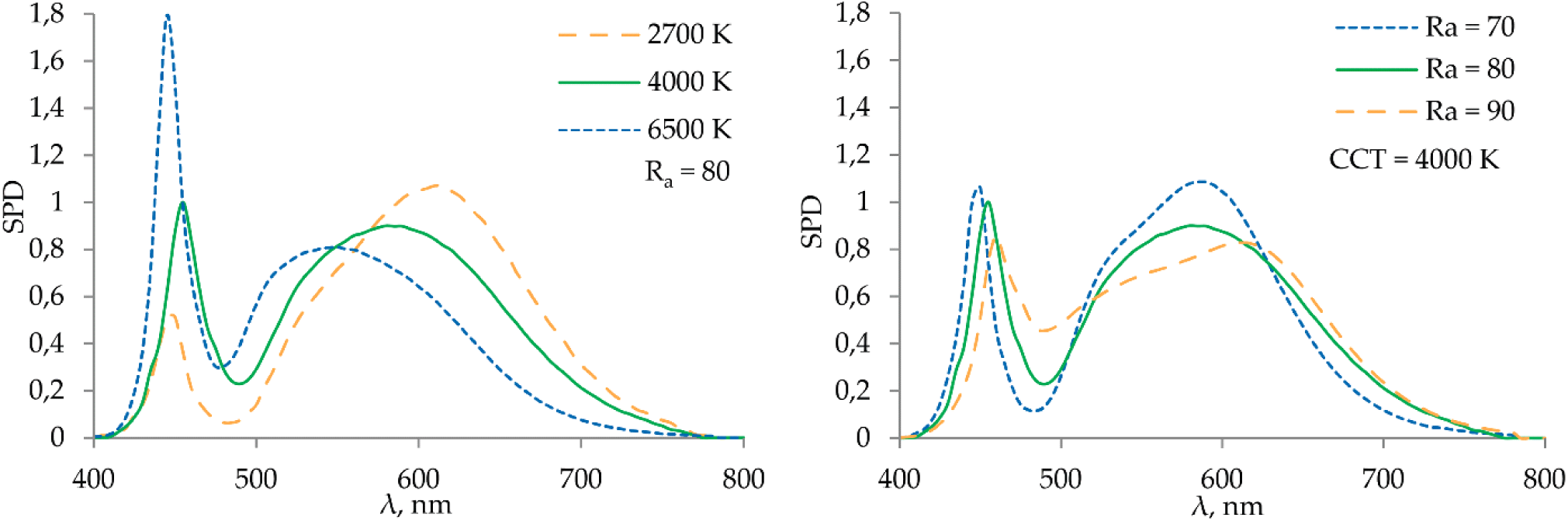
Spectra of white LED light with the same color rendering, but different color temperature CCT (*left*), and with the same color temperature and different color rendering *R*_*a*_ *(right)*

Differences in the spectrum of white diodes with the same color rendering and color temperature are barely perceptible. Therefore, we can estimate the spectro-dependent parameters only by color temperature, color rendering and light efficiency – parameters that are written on the label of a conventional white light lamp.

The results from analysis of the spectra of serial white LEDs are as follows:

1. The spectrum of all white LEDs, even with a low color temperature and with maximum color rendering, as in sodium lamps, has a very small amount of far red light (Fig. 5).

**Fig. 5.**
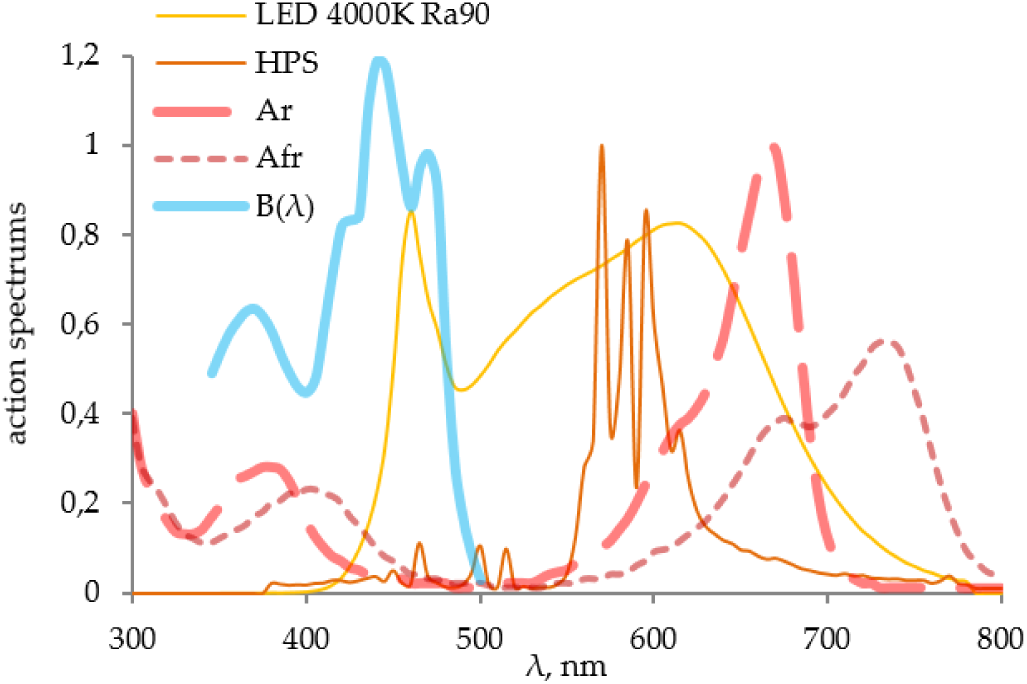
Spectrum of the white LED (*LED* 4000*K R*_*a*_ = 90) and sodium light (*HPS*) in comparison with the spectral functions of the plant susceptibility to blue (*B*), red (*A*_*r*_) and far red light (*A*_*fr*_)

In natural conditions, a plant shaded by the leaves of other plants receives more far red light than near red light, which triggers a ‘shadow avoidance syndrome’ in light-loving plants – a plant grows upward. For example, tomatoes need far red light at the stage of growth (not seedlings!) to stretch, grow bigger, increase a total occupied area, and, consequently, the yield in the future.

Accordingly, under white LEDs and sodium light, a plant feels like under the direct sunlight and does not grow upward.

2. Blue light is needed for the reaction of phototropism (Fig. 6).

**Fig. 6.**
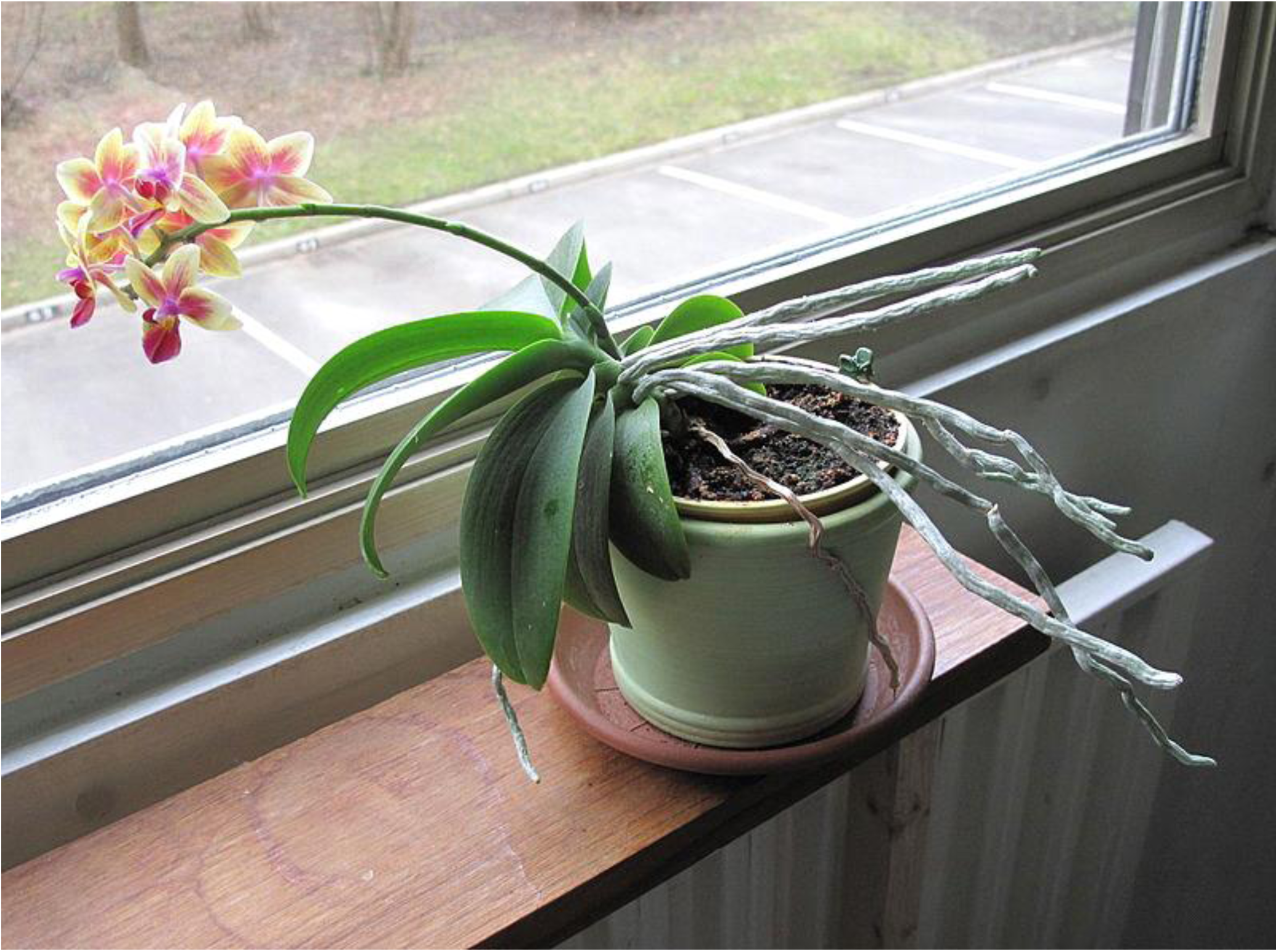
Phototropism – bending of leaves, flowers and stems towards a blue component of white light (photo from Wikipedia)

A phytoactive blue component in one watt of a 2700 K white LED flux is two times bigger than in one watt of sodium light. And the amount of phytoactive blue component in white light grows in proportion to the color temperature. If, for example, decorative flowers need to be moved toward people, they should be illuminated from this side by intense cold light.

3. Energy value of light is determined by the color temperature and color rendering and can be determined by the following formula with an accuracy of 5%:

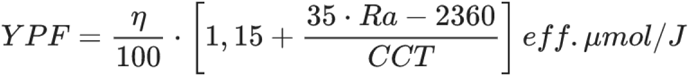

where *η* — light output in *lm / W*, *Ra* — overall color rendering index, *CCT* — correlated color temperature in degrees Kelvin.

Examples of using this formula:

A. Let’s find an illumination value for the basic values of the white light parameters to ensure, for example, 300 eff. μmol / s / m^2^ at given color rendering and color temperature:

| **White LED illumination in kiloluxes corresponding to 300 eff. $\mu\text{mol} / \text{s} / \text{m}^2$** |  |  |  |
| --- | --- | --- | --- |
| **CRI \ CCT** | **3,000 K** | **4,000 K** | **5,000 K** |
| **Ra = 70** | 25 klx | 26 klx | 26 klx |
| **Ra = 80** | 23 klx | 24 klx | 24 klx |
| **Ra = 90** | 21 klx | 22 klx | 23 klx |
| **Ra = 95** | 20 klx | 22 klx | 22 klx |

You can see that the use of warm white light with high color rendering allows using slightly smaller illumination values. But if you consider that the light output of warm LEDs with high color rendering is somewhat lower, it becomes clear that you cannot get any significant energetic advantages or disadvantages by selecting color temperature and color rendering. You can only adjust the proportion of phytoactive blue or red light.

B. Let’s estimate the applicability of a typical general-purpose LED lamp for growing microgreens.

Assume that the lamp with a size of 0.6 × 0.6 m consumes 35 W and has a color temperature of 4,000 K, color rendering *Ra* = 80 and light output of 120 lm / W. Then its efficiency will be *YPF* = (120 / 100) × (1.15 + (35 × 80 – 2,360) / 4000) eff. μmol / J = 1.5 eff. μmol / J. That will be 52.5 eff. μmol / s if you multiply it by 35 W.

If you put this lamp sufficiently low above a bed of microgreens with an area of 0.6 × 0.6 m = 0.36 m^2^ and thereby avoid light loss to the sides, the illumination density will be 52.5 eff. μmol / s / 0,36m^2^ = 145 eff. μmol / s / m^2^. This is about half the usual recommended values. Consequently, the lamp power must also be doubled.

## Direct comparison of phytoparameters for fixtures

Compare the phytoparameters of a conventional office ceiling LED fixture produced in 2016 and specialized phyto lamps (Fig. 7).

**Fig. 7.**
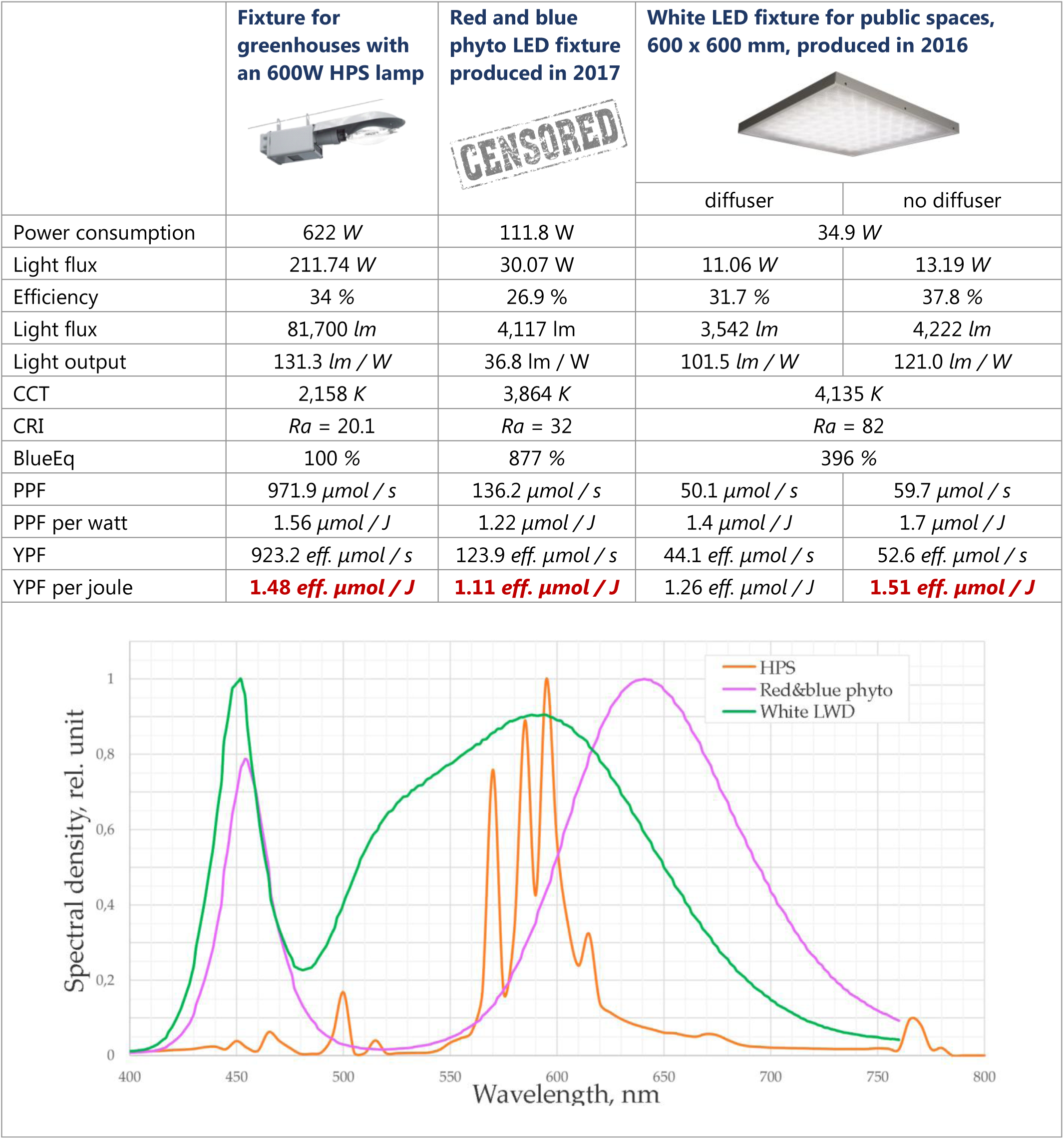
Comparative parameters of a typical 600W sodium lamp for greenhouses, a specialized phyto LED fixture and a general lighting fixture

You can see that a conventional general lighting fixture with a removed diffuser has the same energy efficiency for plant lighting as a specialized sodium lamp. You can also see that the red and blue phyto fixture (the manufacturer is not specified intentionally) has a lower technology level, since its full efficiency (the ratio of the light flux power in watts to the power consumed from the mains) is smaller than the efficiency of the office light fixture. But if the efficiency values of the red and blue and white fixtures were the same, then the phytoparameters would also be approximately the same!

You can also see from the spectra that the red and blue phyto fixture is not narrow-banded, its red hump is wide and contains much more far red light than the white LED fixture and sodium lamp. In those cases where far red light is required, the use of this fixture alone or in combination with other options may be appropriate.

## Estimation of energy efficiency of the lighting system as a whole

The author uses a hand-held spectrometer UPRtek 350N (Figure 8) provided by <u>**Intech Engineering**</u>.

**Fig. 8.**
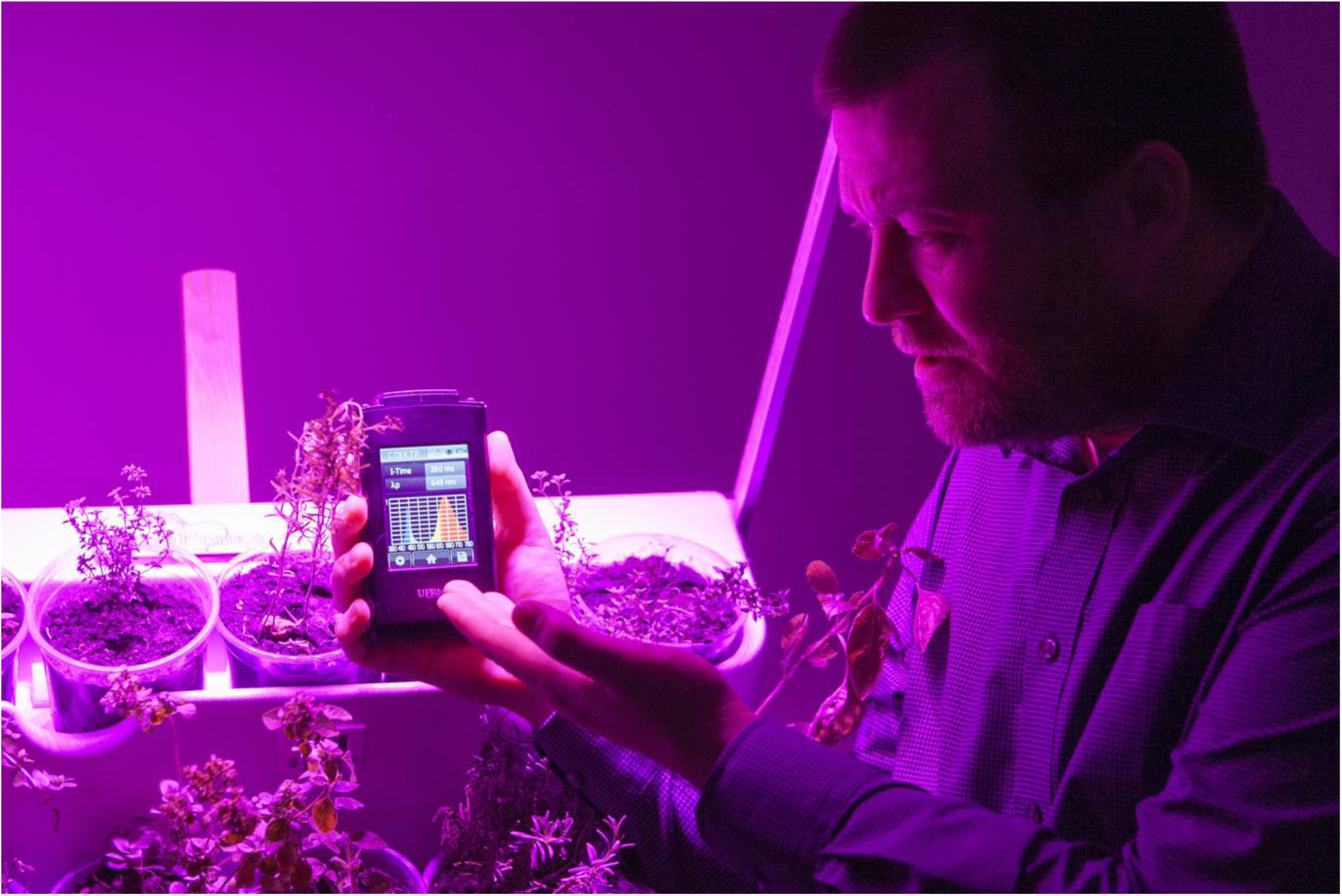
Auditing the phyto lighting system

The next *UPRtek* model is the *PG100N* spectrometer that measures directly micromoles per square meter, effective micromoles per square meter and, more importantly, light flux in watts per square meter.

Measurement of the light flux in watts is an excellent function! If you multiply the illuminated area by the light flux density in watts and compare it with the lamp consumption, you can find energy efficiency of the lighting system. Today, it is the only indisputable criterion of effectiveness, which in practice differs by an order of magnitude for different lighting systems (not by a several-fold factor or even more by percents, as the energy effect changes when the shape of the spectrum is changed).

### Examples of using white light

Examples of using both red-blue and white light for hydroponic farms are described (Fig. 9).

**Fig. 9.**
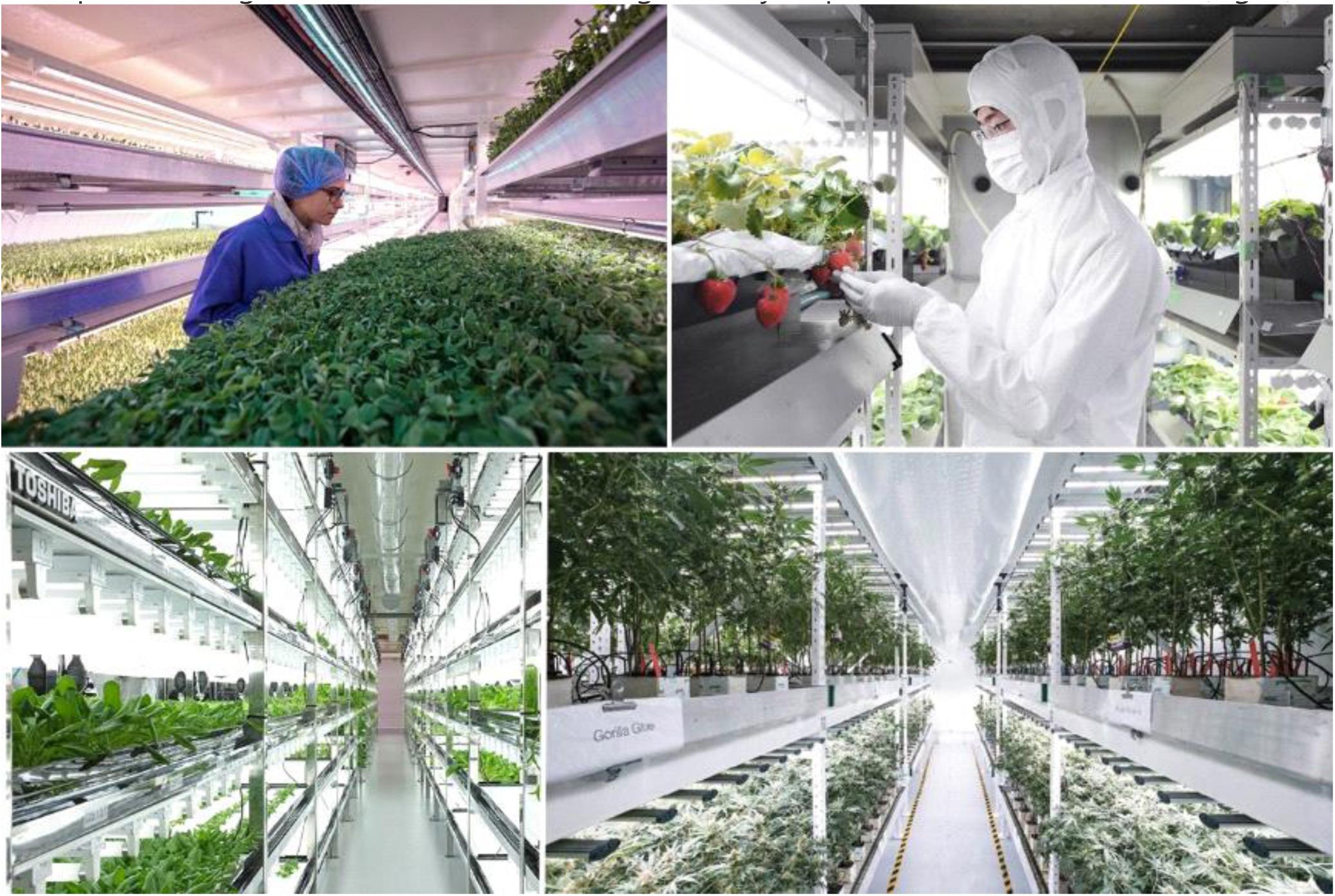
From left to right and from top to bottom: *Fujitsu*, *Sharp*, *Toshiba*, medicinal plant farm in Southern California

A fairly well known system of farms is *Aerofarms* (Fig. 1, 10) with the largest farm built close to New York. In *Aerofarms*, more than 250 kinds of greenery are grown under white LED lamps with more than twenty harvests a year.

**Fig. 10.**
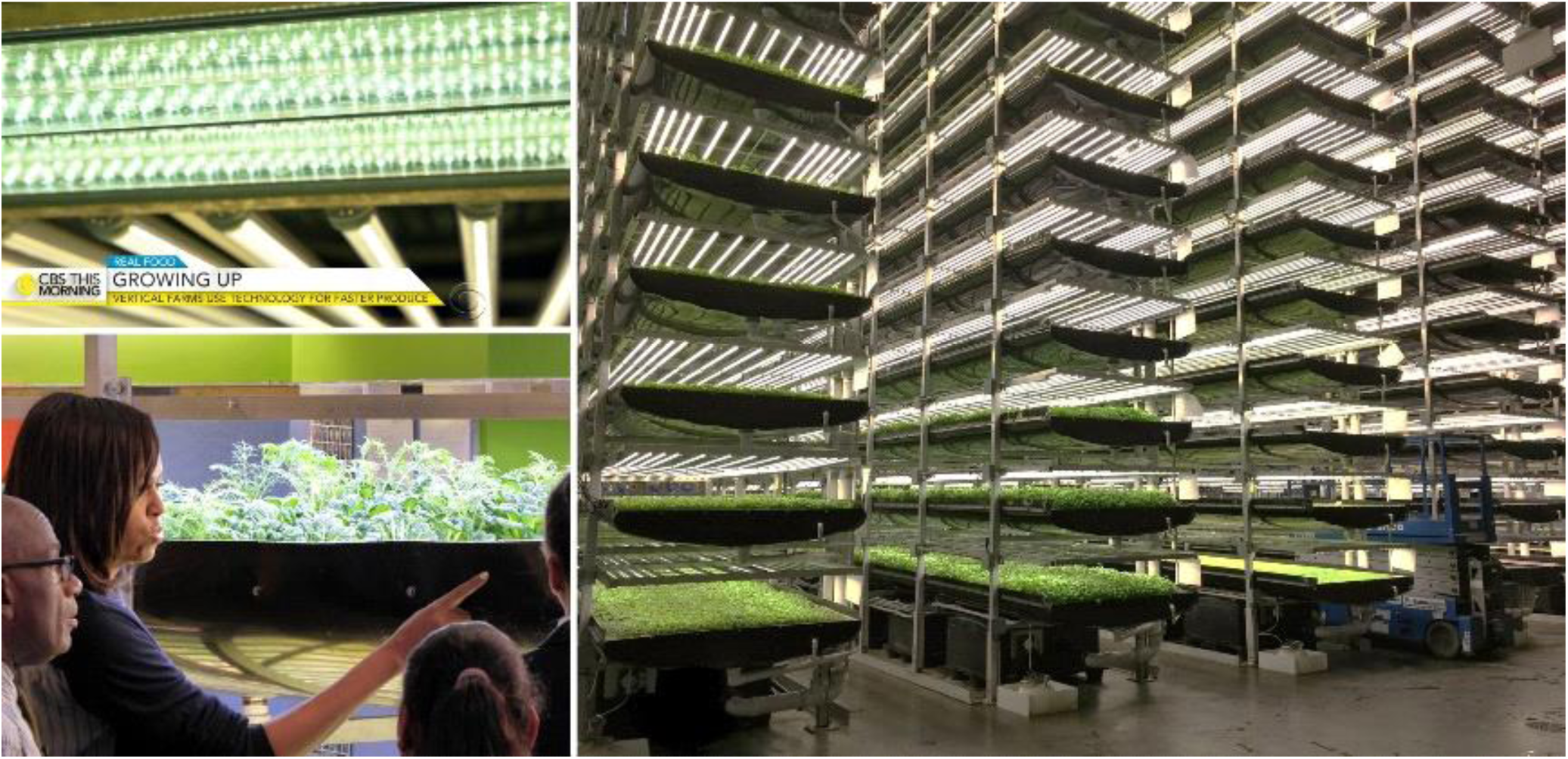
*Aerofarms* in New Jersey (*Garden State*) on the border with New York

## Direct experiments to compare white and red-blue LED lighting

The published results of direct experiments comparing plants grown under white and red-blue LEDs are extremely insufficient. For example, some of these results were shown at the Moscow Timiryazev Agricultural Academy (Fig. 11).

**Fig. 11.**
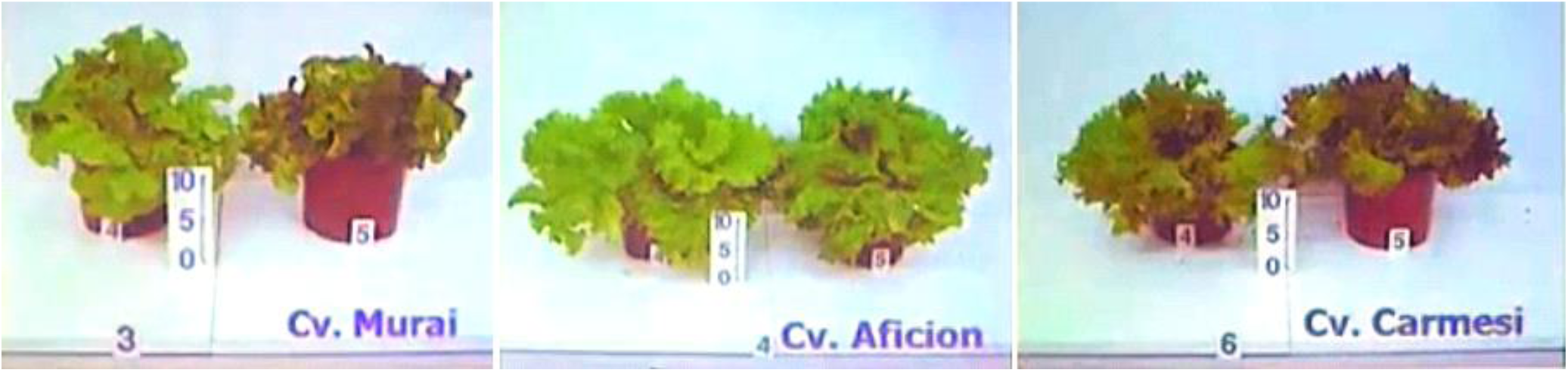
In each pair, the plant on the left is grown under white LEDs, and the plant on the right – under red-blue LEDs (from the <u>**presentation**</u> by I.G. Tarakanov, Department of Plant Physiology, Moscow Timiryazev Agricultural Academy)

In 2014, the Beijing University of Aeronautics and Astronautics published the results of a large study of wheat grown under different types of LEDs [4]. The Chinese researchers concluded that it was advisable to use a combination of white and red light. But if you look at the figures from the article (Fig. 12), you can notice that the difference in parameters for different types of lighting is not dramatic at all.

However, the main trend of today’s research is correcting the shortcomings of narrow-band red and blue lighting by adding white light. For example, Japanese researchers [5, 6] have shown an increase in the mass and nutritional value of lettuce and tomatoes when white light is added to red light. In practice, this means that if the aesthetic appeal of the plant during growth is unimportant, it is not necessary to reject narrow-band red and blue lamps that have already been bought. White light fixtures can be used additionally.

**Fig. 12.**
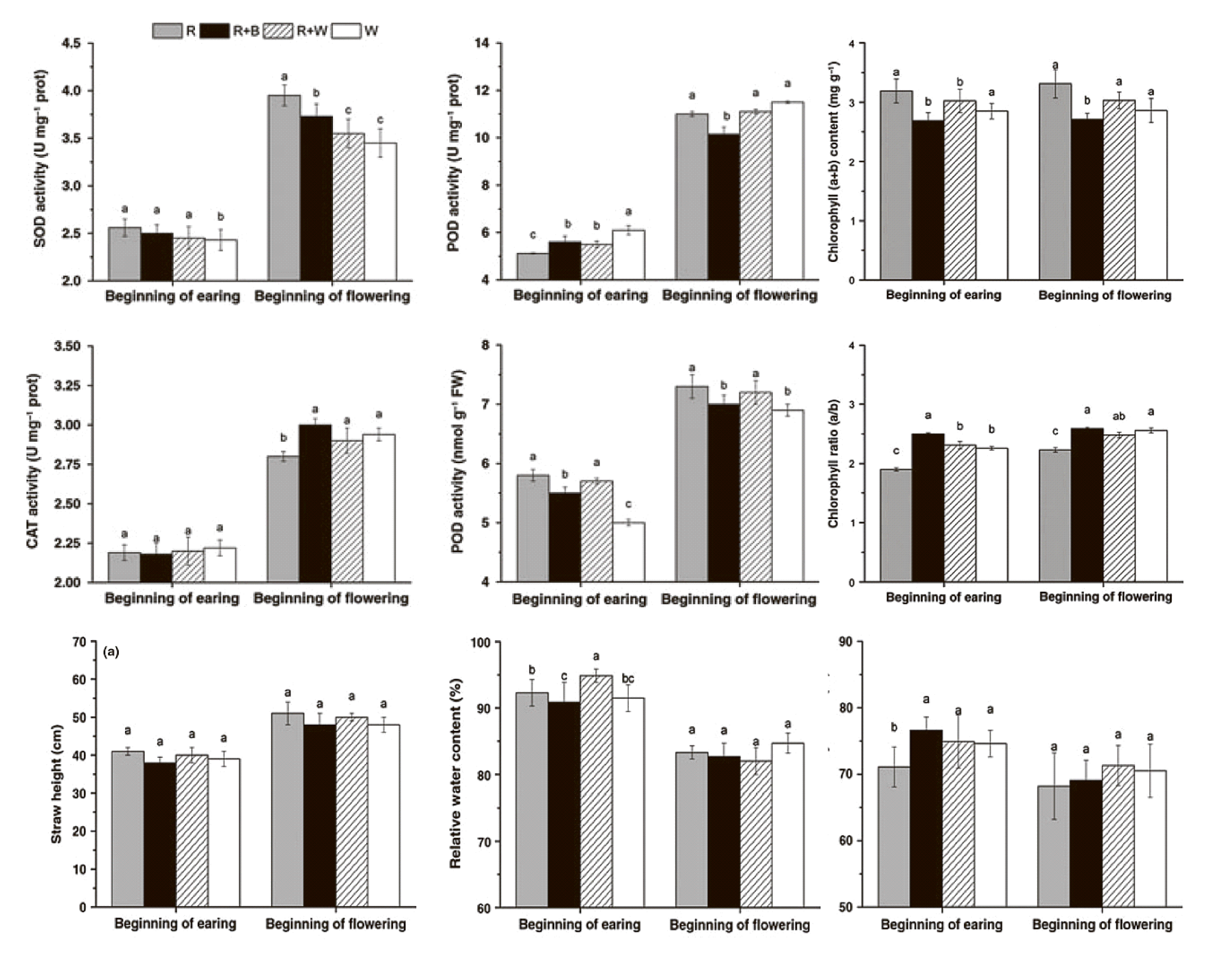
Values of the factors studied in two phases of wheat growth under red, red and blue, red and white, and white LEDs

## Influence of the light quality on the result

The fundamental ecology law, *Liebig’s Law*, (Fig. 13) states that growth is dictated not by total resources available, but by the scarcest resource. For example, if there is no lack of water, minerals and CO2, but the lighting intensity is 30% of the optimal value – a plant will give no more than 30% of the maximum possible yield.

**Fig. 13.**
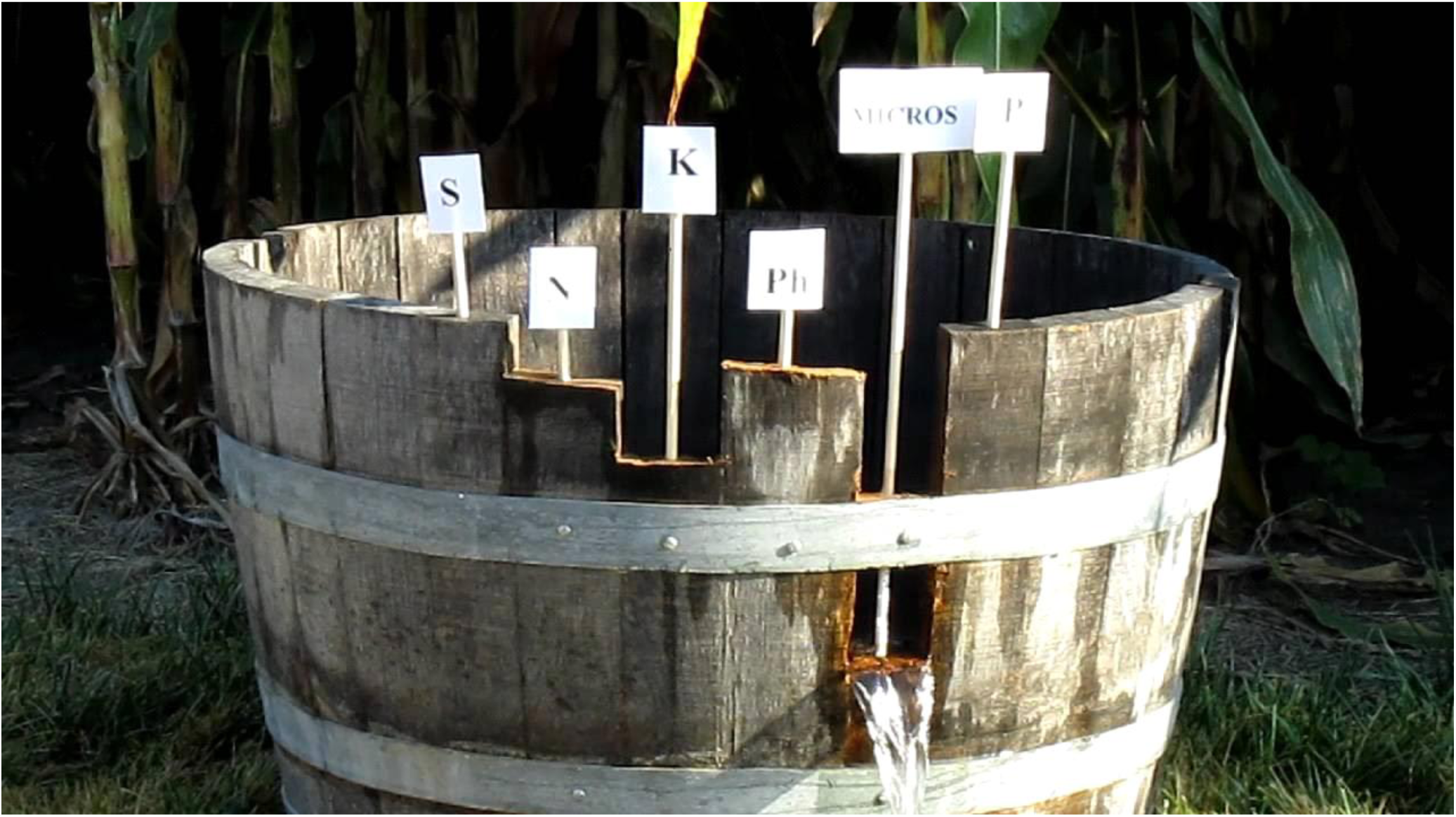
Illustration of the limiting factor principle from the <u>**training video**</u>

The plant’s response to light – intensity of gas exchange, consumption of nutrients from the solution and synthesis processes – is determined by laboratory methods. Responses characterize not only photosynthesis, but also the processes of growth, flowering, and synthesis of substances necessary for taste and aroma.

Fig. 14 shows the plant’s reaction to the change in the wavelength of lighting. The authors measured the intensity of consumption of sodium and phosphorus from the nutrient solution by mint, strawberry and lettuce. Peaks on these graphs are signs of stimulating a specific chemical reaction. The graphs show that excluding some ranges from the full spectrum for the sake of economy is like removing some of the piano keys and playing a melody using the rest ones.

**Fig. 14.**
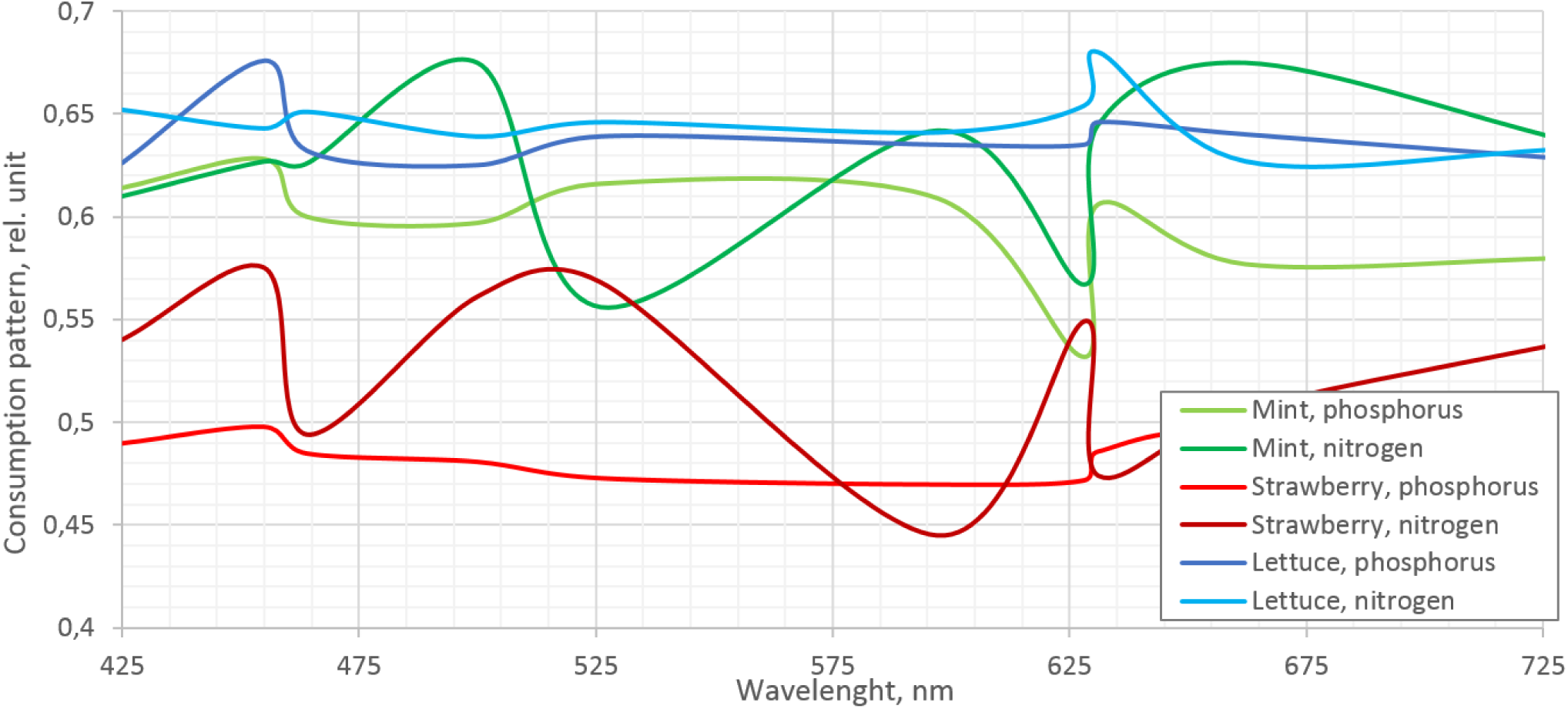
Stimulating role of light for nitrogen and phosphorus consumption by mint, strawberry and lettuce (data provided by <u>**Fitex**</u>)

The limiting factor principle can be extended to separate spectral components – in any case, a full spectrum is needed for a full-fledged result. Removing certain ranges from the full spectrum does not lead to a significant increase in energy efficiency. Moreover, the Liebig’s Law can occur and the result will turn out to be negative.

The examples demonstrate that the usual white LED light and special ‘red and blue phyto light’ for plants have about the same energy efficiency. But the broadband white light comprehensively satisfies the plant’s needs expressed not only in the stimulation of photosynthesis.

Removing green light from the continuous spectrum so that white light turns into purple is a marketing trick for buyers who want a ‘special solution’, but are not qualified customers.

## White light correction

The most common general-purpose white LEDs have low color rendering *Ra* = 80, which is caused by a lack of red color in the first place (Fig. 4).

The lack of red light in the spectrum can be replenished by adding red LEDs to the lamp. This solution is promoted, for example, by <u>**CREE**</u>. The Liebig’s law logic suggests that this adding will do no harm, if it really is an adding of red LEDs, and not redistribution of energy from other ranges in favor of red light.

An interesting and important work was carried out in 2013-2016 at the Institute of Biomedical Problems [7, 8, 9]: researchers investigated how adding narrow-band red LEDs of 660 nm to white LEDs of 4,000 *K / Ra* = 70 affect the growth of Chinese cabbage.

The results were the following:

- Under the LED light, the cabbage growth is almost the same as under a sodium lamp, but cabbage has more chlorophyll (its leaves are greener).
- The dry yield weight is almost proportional to the total amount of light in moles received by the plant. The more light, the bigger yield of cabbage.
- The concentration of vitamin C in cabbage slightly grows as the lighting increases, but it becomes significantly bigger when red light is added to white light.
- A significant increase in the proportion of the red component in the spectrum increased the concentration of nitrates in biomass dramatically. The researchers had to optimize the nutrient solution and introduce part of nitrogen in the ammonium form, so as not to exceed the MAC for nitrates. But under pure white light it was possible to work with a nitrate form only.
- At the same time, an increase in the proportion of red light in the total light flux has almost no effect on the yield mass. That is, replenishment of the missing spectral components affects not the amount of yield, but its quality.
- Higher efficiency of a red LED in moles per watt makes adding red light to white light effective energetically.

Thus, the addition of red light to white light is advisable in the particular case of Chinese cabbage and is quite possible in the general case. Of course, it should be done under biochemical control and with proper selection of fertilizers for a particular crop.

## Options for the spectrum enrichment with red light

A plant does not know where white or red light spectrum quanta come from. There is no need to make a special spectrum in one LED. And there is no need to use red and white light from one special phyto fixture. It is sufficient to use general-purpose white light and take a separate red light lamp to illuminate the plant additionally. And when a person is near the plant, a red lamp can be turned off using the motion sensor, so that the plant looks green and pretty.

But a reverse solution is also justified – by selecting the luminophore composition, you can expand the white LED emission spectrum towards long waves and balance it so that the light remains white. And you will get white light with extra-high color rendering suitable both for plants and humans.

It is especially interesting to increase the share of red light by increasing the overall color rendering index for <u>**city farming**</u> – a social movement that involve growing the plants necessary for man in the urban areas, often with people and plants sharing their living space and, consequently, light environment.

## Open questions

We can try to identify the role of the ratio between far and near red light and the advisability of using a ‘shadow avoidance syndrome’ for different cultures. We can argue in which areas we should break the scale of wavelengths for the analysis.

We can discuss whether a plant requires wavelengths shorter than 400 nm or longer than 700 nm for stimulation or a regulatory function. For example, there is a private message that ultraviolet significantly affects the consuming properties of plants. Among other things, the red-leaved types of lettuce are grown without ultraviolet, and they grow green, but before the sale they get irradiated with ultraviolet. After that they turn red and are taken for sale. And how well the new *PBAR* metrics (*plant biologically active radiation*) described in the standard *ANSI/ASABE S640*, *Quantities and Units of Electromagnetic Radiation for Plants (Photosynthetic Organisms*) prescribes to consider a range of 280-800 nm.

## Conclusion

Network stores choose more long-keeping types, and then buyers vote with their money for more vivid fruits. And almost no one chooses taste and aroma. But as soon as we become richer and begin to demand more, science will instantly give the necessary types and recipes of nutrient solution.

To make a plant synthesize everything that is needed for taste and aroma, you need lighting with a spectrum containing all wavelengths to which the plant will react, i.e. in the general case, a continuous spectrum. Perhaps the basic solution will be white light with high color rendering.

## Acknowledgement

The author expresses sincere gratitude for the help in the preparation of the article to Irina Konovalova, PhD in biology at the Institute of Biomedical Problems; Tatiana Dubovskaya, the head of the city farming school <u>**UrbaniEco**</u>; Tatiana Trishina, the head of the <u>**Fitex project**</u>; and Mikhail Chervinskiy, expert at CREE.

